# Representational geometry of emotional empathy in the ventromedial prefrontal cortex and in the mid-posterior insula

**DOI:** 10.1101/2025.08.30.673208

**Authors:** Leonardo Cerliani, Rune Bruls, Cas Teurlings, Settimio Ziccarelli, Valeria Gazzola, Christian Keysers

**Author notes:** Equal contribution (Valeria Gazzola, Christian Keysers).

## Abstract

Facial expressions provide rapid and informative cues about others’ emotional and mental states, playing a critical role in social interactions. However, whether distinct emotional expressions reflect discrete neural processes or arise from varying combinations of underlying affective dimensions such as arousal and valence remains a subject of investigation.

Crucially, these accounts need not be mutually exclusive: different brain regions may encode emotional expressions along both categorical and dimensional axes to varying degrees. To test this hypothesis, we probed different brain systems involved in emotion recognition - the ventral attentional network and the cortical limbic system centered on the ventromedial prefrontal cortex (vmPFC) - to investigate the extent to which these networks and their subregions encode emotional facial expressions in terms of (1) perceived arousal, (2) arousal+valence, or (3) six discrete emotion categories (anger, disgust, fear, happiness, pain, sadness).

To this aim, we modelled the fMRI signal from perceiving movies of facial emotional expressions with ratings for either arousal, arousal+valence or emotion category using representational similarity analysis (RSA) - a method that aims at assessing which ratings model better represents how the perception of facial expressions is reflected by the fMRI parameter estimates across all the voxels within a brain region.

This analysis showed that regions in the vmPFC network, including the subgenual cingulate and the medial OFC, are sensitive to ratings for distinct emotion category and for arousal+valence - significantly more so for the former - while they fail to show sensitivity for arousal ratings alone. In the ventral attentional network, the mid-posterior insula showed a similar profile, while the most posterior and ventral insular show evidence of encoding arousal+valence, but not for emotion category or arousal alone.

Our findings support the idea that the vmPFC and the mid-posterior insula play a role in encoding emotion-specific and valence representations beyond general arousal processing. These results contribute to understanding how integrative brain regions support emotional empathy and social cognition.

## Introduction

The ability to accurately identify our interlocutors’ emotional state from their facial expressions plays a crucial role in shaping how the interaction unfolds. Understanding how our brain enables us to distinguish different emotions is therefore important to understand how it supports social cognition (Adolphs 1999; van de Riet, Grezes, and de Gelder 2009; Barrett et al. 2019).

Reading other people’s emotions from their facial expression results from a complex - albeit highly efficient - integration of interoceptive, cognitive, sensory, and mnestic representations (Barrett et al. 2007). How such integration is carried out in the brain is still a matter of discussion (Pessoa 2008; Gu et al. 2019). Two influential theories have been proposed. Proponents of the basic emotions theory (P. Ekman et al. 1992; J. Panksepp 1998; Izard 2009; Celeghin et al. 2017; Keltner et al. 2019) suggest that we evolved to experience, express and perceive distinct categories of emotions, including at least the 6 basic emotions of anger, fear, disgust, happiness, sadness, and surprise (Paul Ekman and Cordaro 2011; Jaak Panksepp and Watt 2011). In its strongest form, these basic emotions are thought to reflect distinct physiological mechanisms including specialized brain circuits evolved to serve specific purposes. On the other hand, proponents of constructivism (Barrett and Wager 2006; Barrett 2017; Wager et al. 2015; Kober et al. 2008; Lindquist et al. 2012) suggest that identifying specific emotions requires integrating fundamental neurophysiological responses to perceived arousal and valence (core affect) with past experiences and interpretations of the current situation based on high-order social cognitive representations.

From a cognitive neuroscience perspective, these discussions raise the question of which dimensions are encoded in the neural representation of observed facial expression in different brain regions and networks. An investigation aimed at answering these question does not entail arbitrating between basic emotion and constructivist theories, as both acknowledge the presence of high-level cognitive representation for distinct category of emotions. Instead, it speaks to a more mechanistic question of which brain regions show evidence of processing emotional facial expressions in terms of (1) perceived arousal, (2) perceived arousal *and* valence or (3) categorical emotions.

In the present study, we tackle these questions by conjointly analysing fMRI-based brain activity recorded during the observation of videos displaying emotional facial expressions, as well as the ratings of these videos provided by the same participants from the fMRI acquisition. In these videos, four different actors displayed six different emotions (anger, disgust, fear, happiness, pain, sadness), each at two levels of intensity. To estimate which dimensions of facial emotion perception are encoded in different brain regions, we first construct three models representing the variability in behavioural ratings across all movies. The three models include ratings for (1) arousal alone, (2) arousal+valence (hereafter ‘aroval’) or (3) emotion content across six emotions. We then carry out a Representational Similarity Analysis (RSA - (Kriegeskorte, Mur, and Bandettini 2008)) to provide evidence of which behavioural model best fits the variability in fMRI activity across different brain regions.

Rather than carrying out an exploratory analysis on the whole brain, we adopt a hypothesis-driven approach for selecting the brain regions we investigate. Neuroimaging studies have shown that observing emotional facial expressions elicits activity in multiple brain regions, including the fusiform face area (FFA), amygdala, posterior superior temporal sulcus (pSTS), inferior frontal gyrus (IFG), insular and cingulate cortex, as well as the orbitofrontal cortex and medial prefrontal cortex (mPFC) (Haxby, Hoffman, and Gobbini 2000; Adolphs 2001; Zinchenko, Yaple, and Arsalidou 2018). However, compelling evidence suggests that neural signatures of distinct emotions are rarely confined to single regions; rather, they are better predicted by distributed patterns of activity across networks (Chang et al. 2015; Saarimäki et al. 2016). Notably, the ventromedial prefrontal cortex (vmPFC) and insula have emerged as core hubs within these networks (Lindquist et al. 2012; Wager et al. 2015; Roy, Shohamy, and Wager 2012; Seeley et al. 2007).

Therefore, we focused our investigation on two large brain networks derived from a widely used resting-state fMRI parcellation on a large sample of participants (Yeo et al. 2011). Specifically, we examined (1) a limbic network including the orbitofrontal cortex (OFC), the temporal pole, the subgenual cingulate (sgACC) and the ventromedial prefrontal cortex (vmPFC); (2) the ventral attentional network, encompassing regions of the temporoparietal junction, supramarginal gyrus, frontal operculum, and anterior insula. Importantly, since RSA - like other multi-voxel techniques - is heavily dependent on the definition of the region of interest (ROI), we investigated the subdivisions of cortical limbic and ventral attentional network (Corbetta and Shulman 2002; Corbetta, Patel, and Shulman 2008) across different atlases (Yeo et al. 2011; Makris et al. 2006; Eickhoff et al. 2005), each defined using different criteria, in order to provide a robust support to our findings beyond a single parcellation scheme.

## Materials and Methods

### Participants

Twenty-six healthy participants (mean age = 23.04; SD = 2.29; 12 females), took part in the study. All of them were right-handed, had normal or corrected to normal vision, spoke English fluently, were not taking any psychoactive medications that could affect cognitive function, and had no history of neurological or psychiatric illness. The study was approved by the University of Amsterdam (UvA; Protocol 2019-EXT-11148) Ethics Review Board of the Faculty of Social and Behavioral Sciences and was conducted in accordance with the Declaration of Helsinki. Participants were reimbursed for their participation with 10€/hour.

### Stimuli

Participants observed short videos in which four different actors (females aged 29, 27; males aged: 24, 54) each displayed anger, disgust, fear, happiness, pain and sadness at two levels of intensity: high and low. Additionally, each actor displayed a neutral facial expression, devoid of any emotional content - for instance blinking and mouth opening/closing - also at two levels of intensity (here defined by the amount of facial motion). Thus we presented in total 56 movies: 6 movies with emotions + 1 neutral, each at 2 levels of intensity, for each of the 4 actors.

### Experimental Procedure

Participants first took part in the MRI image acquisition during which they were presented the 56 movies in 8 separate and consecutive runs. The same 56 movies were presented in each run, and their occurrence was independently randomized for each run and participants, therefore each participant saw each movie 8 times, one for each run. After the fMRI session, participants completed a behavioural rating task on the same movies.

In the scanner, participants were instructed to observe the movies while feeling with the actor, at the same time trying to avoid deliberately mimicking the facial movements and verbalizing the content of the video. Videos lasted 1.5 s and were separated by a fixation cross with a random duration between 3-10 s. In total, the whole session (8 fmri runs + additional MRI scans) took approximately 75 minutes.

In the subsequent behavioural session, participants were again presented with the same movies and asked to rate each movie along different dimensions. Participants were asked to rate each video on a scale from 0 (min) to 10 (max) along seven dimensions, namely amount of arousal and amount of specific emotional content for each of the 6 emotions. The same rating task was also carried out for a final eigth dimension: the degree of negative or positive valence in the video. In this case the scale range was from -5 to 5. To minimize the effect of rating a video on one dimension on the rating on other dimensions, we used a method similar to that of Adolphs et al. (Adolphs et al. 2000) and presented each video 8 times, asking the participant to rate only one dimension for each presentation. This meant that a participant would need to rate a video of a face expressing Pain also on Anger, etc.. Specifically participants were asked to rate each video on: ‘arousal’, as the ‘state of being bodily alert and awake’, using the question ‘How aroused you perceive the person in the video to be?’ from 0 (min) to 10 (max); ‘valence’ as ‘How positive or negative the state of the person in the video is?’ from -5 (totally negative) to 5 (totally positive); Each of our 6 target emotions as ‘how intense you perceive [sadness, anger, fear, disgust, pain, happiness] to be [in the video]’ on a scale from 0 (min) to 10 (max).

### MRI Image Acquisition and Preprocessing

#### Image Acquisition

All MRI images were acquired on a Philips 7T Achieva at the Spinoza Center in Amsterdam using a 32-channel receive and 2-channel transmit volume coil (Nova Medical, USA).

*fMRI acquisition:* 3D FFE EPI sequence at 1.6 mm isotropic resolution, FOV 205 x 176, 110 slices. Each 3D volume was acquired in 1.82 s, EPI factor = 47, SENSE(AP) = 2.73, flip angle 13°, TE = 19.45 ms, TR = 1816 ms. 8 runs per participant, with 275 volumes per run. For each run, we acquired additional 5 volumes with identical acquisition parameters except for the reversed phase encoding direction (PA), to be subsequently used to correct for distortions.

*anatomical T1w acquisition*: MPRAGE at 0.8 mm isotropic resolution, FOV 288 x 288, 232 sagittal slices, flip angle 8°, TE = 3.29 ms, TR = 3000 ms.

The PAR/REC stacks exported from the Philips scanner were converted to NIFTI images using dcm2niix v1.0.20201102 (https://github.com/rordenlab/dcm2niix) and reoriented to the standard RPI coordinate system using fslorient2std, to enable inspecting the images in a familiar orientation.

#### Image Preprocessing

The scripts used to carry out image preprocessing are available in the github repo at the link: https://github.com/leonardocerliani/RSA/tree/main/prep_scripts [subject to modification until publication, *ndr*]. FMRI data processing was carried out using FEAT (FMRI Expert Analysis Tool) Version 6.00, part of FSL (FMRIB’s Software Library, www.fmrib.ox.ac.uk/fsl) as well as ANTs (https://stnava.github.io/ANTs/) (Avants et al. 2011). The following preprocessing steps were carried out:

- T1w images: skull stripping using antsBrainExtraction and the Oasis template as a reference. Of note, antsBrainExtraction runs the N4 algorithm (using the N4BiasFieldCorrection ANTs command (Tustison et al. 2010)) in order to mitigate the effect of field inhomogeneities which can potentially affect the quality of the brain/skull segmentation. Results were inspected on a subject basis.
- fMRI susceptibility-induced distortion correction: FSL topup (Jesper L. R. Andersson, Skare, and Ashburner 2003) using epi images with the same parameters of the functional acquisition and reverse phase encoding along the anterior-posterior axis
- The 4D images output by topup were further preprocessed in preparation for GLM analysis (carried out in the native fmri space) and registration with T1w images. Specifically, the following pre-statistics processing was applied: motion correction using MCFLIRT (Smith 2002), non-brain removal using BET (Mark Jenkinson et al. 2002), spatial smoothing using a Gaussian kernel of FWHM 5mm, grand-mean intensity normalisation of the entire 4D dataset by a single multiplicative factor, high-pass temporal filtering (Gaussian-weighted least-squares straight line fitting, with sigma=30.0s)
- Registration of the fmri imges to the T1w image was carried out using the BBR algorithm (Greve and Fischl 2009) in FLIRT (M. Jenkinson and Smith 2001; Mark Jenkinson et al. 2002). Registration from T1w to standard space (MNI152 2mm resolution) was initiated with FLIRT and further refined using FNIRT nonlinear registration (J. L. R. Andersson, Jenkinson, and Smith 2007) using both the skull-stripped and original (N4 bias corrected) T1w image. The outcome of registration was inspected and assessed using the report generated by FSL FLIRT/FNIRT. A composite fmri-to-MNI transformation was then generated to limit the amount of interpolations needed to bring the subject-level images in the MNI template. For the same reason, this transformation was applied to the beta maps derived from the subject/run-level fMRI analysis, while the latter was carried out in the native acquisition space.

### Behavioural Analysis

The analysis of behavioural rating was designed to provide evidence of the validity of the subsequent RSA analysis. Therefore we examined both how much the participants were able to correctly identify the main emotion displayed in each movie, as well as whether movies representing different emotions would elicit substantially different levels of arousal and valence.

The congruence between the displayed and highest rated emotion allowed us to assess the extent to which participants were engaging in brain activity related to that specific emotion, while establishing that movies of different emotion elicit clearly separated levels of arousal and valence was crucial to provide the source of variability between movies which will be then related to the corresponding brain activity during RSA analysis.

To assess whether participants correctly identified the emotion displayed in each movie - hereafter referred to as “congruence” between the displayed and the highest rated emotion in the emotion rating task - we estimated how typical was the emotion rating of a given participant and each movie by comparing her rating (a vector of 6 values) to the average rating of all other participants using Pearson correlation. This allowed us to assess whether and which participant carried out the emotion rating task according to expectations - providing a clearly highest rating for the emotion which is actually displayed in the movie. Being confident that participants correctly identified most of the emotions displayed in the movies provides a crucial element for the interpretation of the fMRI results and especially of the RSA analysis.

In order to assess whether different movies elicited different ratings in terms of Arousal and Valence, we carried out two separate repeated-measurement analysis of variance (RM-ANOVA), with factors for Emotion (6 levels, corresponding to the emotion being displayed in each movie) and Intensity (of the displayed emotion: high or low). Subsequent post-hoc tests were corrected for multiple comparisons with Holm correction.

### Representational Similarity Analysis (RSA)

#### RSA Motivation and Overview

We aimed at assessing in which brain regions the fMRI activity elicited by other people’s emotional facial expression reflect a more categorical representation of others’ emotions corresponding to six different emotion types, or rather encodes the amount of perceived arousal, or the combination of perceived arousal and valence.

This investigation could be carried out using a mass-univariate approach highlighting voxels that show any difference in the modulation of the brain signal due to rating for either emotional content, arousal or arousal+valence (hereafter: ‘aroval’). However, the identification of a specific emotional state during social interaction rests on cognitive functions which integrate cognitive, sensory and interoceptive neural representations of currently unfolding events, previous experiences, social norms and future plans. Such cognitive integration is supported by higher-order brain regions where precise localization of cognitive function is difficult at the level of single voxels, and where instead functional specialization can results from a complex encoding of stimuli/tasks across all voxels within the brain region, rather than being reflected in the signal of specific voxels, or in their average activity.

Accordingly, in addition to the voxelwise parametric modulation analysis (the results of which are available in the Supplementary Materials), we implemented a Representational Similarity Analysis (RSA, (Kriegeskorte, Mur, and Bandettini 2008)), which retains as its initial source of information the whole pattern of activity in response to a specific stimulus across all voxels in a given brain region - i.e. a vector of *n* beta values from the voxelwise analysis, where *n* is the number of voxels in that brain region. Computing this vector for *m* movies yields an *m-by-n* matrix which represents the brain activity for each voxel of that brain region across different movies. Subsequently, calculating the (dis)similarity between each and each other movie’s vector yields a symmetric *n-by-n* matrix - hereafter called representation dissimilarity matrix (RDM).

This RDM provides us a way to represent the differences in the response to different emotion movies across all the voxels in a brain region. To relate these differences in brain activity to differences in ratings - emotional content, arousal, arousal + valence - we can repeat the calculation of the RDM for the ratings themselves. These RDMs for behavioural ratings will represent the *models* we will fit to the RDM of brain activity to identify brain regions where the differences in stimuli encoding are related to the cognitive processes of rating those movies. In its simplest form, this is carried out by estimating the correlation coefficient between the unique <u>^𝑛(𝑛−1)^</u> elements (upper or lower triangular, after removing the uninformative diagonal) of either RDMs. The resulting RSA score therefore represents the similarity/correlation between the fMRI and the ratings RDM.

At this point we can test in which regions the mean RSA scores are significantly different from zero across participants. Finally, we can compare the RSA scores for different models, to identify which regions appear to be more specialized for encoding either categorical emotions, arousal or arousal+valence.

#### RSA Regions of interest

One further element to consider is the choice of the regions of interest to run RSA on. We are interested in brain regions sensitive to differences in either the type of emotion, or to the associated arousal and valence. This led us to focus our investigation on two networks of interest: (1) a limbic network including the orbitofrontal cortex (OFC), the temporal pole, the subgenual cingulate (sgACC) and ventromedial prefrontal cortex (vmPFC); (2) the ventral attentional network, encompassing regions of the temporoparietal junction, supramarginal gyrus, frontal operculum, and anterior insula. Both were selected from the initial 7-networks parcellation of (Yeo et al. 2011) based on resting-state fMRI data (Fig. 1).

**Fig. 1.**
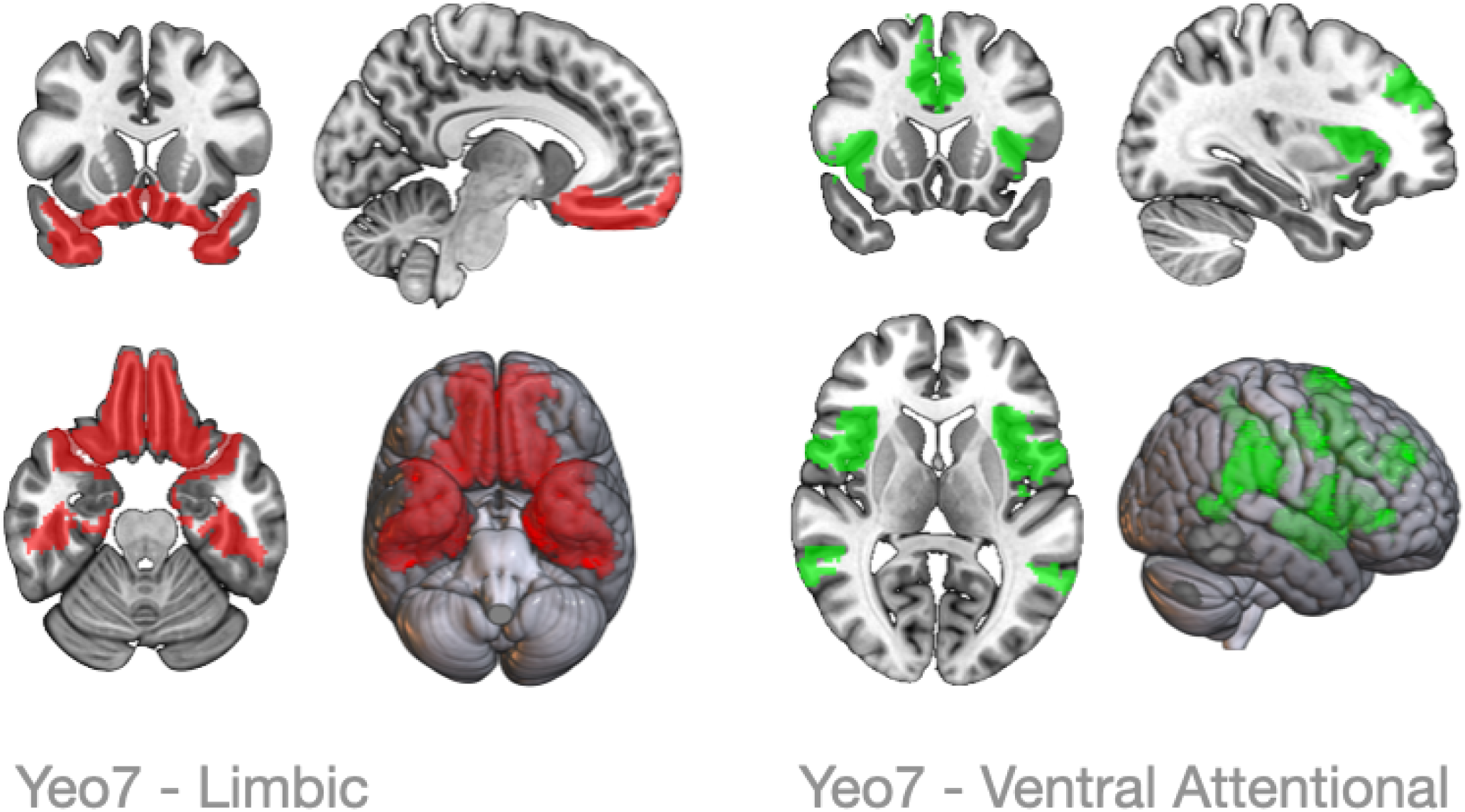
: Overlay of the two large-scale networks obtained from resting-state fMRI in (Yeo et al. 2011). The Limbic network (Yeo7 ROI #5) includes the orbitofrontal cortex (OFC), the temporal poles (TP) and the anterior parahyppocampal gyrus. The Ventral Attentional network (Yeo7 ROI #4) includes regions in the posterior parietal and temporal cortex (supramarginal gyrus, temporo-parieteal junction TPJ), the frontal operculum and the adjacent ventro-lateral premotor cortex (vlPMC), the mid-cingulate and adjacend mid-line premotor regions, part of the middle frontal gyrus as well as the whole anterior and middle insula, excluding only the most dorsal and posterior part adjacent to the Heschl gyrus.

We decided to focus our RSA analysis on these two brain networks because they encompass many regions which have been consistently found to be involved in emotion discrimination and arousal/valence of the emotional content. At the same time, since the outcome of RSA analysis is heavily dependent on the spatial definition of the ROI, we also investigated further subdivisions of these networks, both with the Yeo17 network parcellation, as well as with regions from alternative parcellation schemes (Makris et al. 2006; Eickhoff et al. 2005). This hierarchical and multi-atlas RSA analysis was carried on in order to (1) further add to the evidence found with one specific parcellation; (2) to gain better spatial resolution on the potential sub-regions involved in a given RSA results; (3) and to limit the possibility of false negatives due to having chosen too narrow or too large ROIs.

#### RSA implementation

RSA scores were calculated using Pearson correlation coefficient of the unique elements of the two representation similarity matrices derived from fMRI brain activity and from ratings.

In both cases, the fmri beta values and ratings associated with the same emotion movie across 4 different actors were averaged, to obtain in total 12 data points for each participant, corresponding to 6 data points for movies displaying the same emotion in high intensity (anger, disgust, fear, happiness, pain, sadness) and 6 data points for movies displaying the same emotion in low intensity. The fmri and ratings RDMs were constructed as follows:

**In the behavioural (ratings) domain**, the RDMs were calculated for each participant starting from the matrices of *m* = 12 data points by *k* dimensions, where *k*=1 for arousal rating and *k*=2 for arousal+valence (aroval) rating and *k*=6 for the emotion rating. We calculated their distance matrix using Euclidean distance (cosine similarity would have been uninformative for scalars, while Mahalanobis distance for scalars reduces to Euclidean distance).

**In the fMRI domain**, the RDMs were calculated for each participant starting from an *m-by-v* matrix where *m* is again the number of data points (*m* = 12) and *v* the number of voxels within a specific ROI. The values in each cell are the parameter estimates (betas) obtained from the GLM carried out with FSL Feat. Specifically, these betas were obtained on a subject-level for each of the 8 runs, in a GLM model with one predictor for each stimulus. This model also included the temporal derivatives of the predictors, as well as movement parameters (estimated by FSL mcflirt) and their first derivative. A higher level FSL Feat modeled the average across the 8 runs, and resulted in the beta values eventually used to calculate the distance matrices between movies across all voxels in an ROI. For coherence with the similarity estimates in the behavioural data - and following common practices - Euclidean distance was chosen for fMRI data as well.

This procedure yielded 4 symmetric RDMs: one derived from fMRI beta values, the other three from each model: arousal, aroval and emotion content ratings, respectively. RSA scores were then calculated by testing against zero the correlation of the unique elements (upper or lower triangular without diagonal) of the fMRI RDM with the unique elements of each rating type (after Fisher r-to-z transformation). To establish whether a model would significantly explain more variance in the fMRI RDMs than the other two models, we used a two-tailed paired t-test and applied correction to correct for multiple comparisons (6 pairwise comparisons in total) using false discovery rate.

This procedure was carried out first for the initial two Yeo7 ROIs - limbic and ventral attentional network - and subsequently for different subset of voxels according to the subdivisions of Yeo17, the Harvard-Oxford cortical atlas (Makris et al. 2006), as well as the Anatomy Toolbox (https://github.com/inm7/jubrain-anatomy-toolbox) (Zilles and Amunts 2010).

As an additional precautionary analysis, we also carried out RSA after removing motion energy from the movies (quantified as the sum of the variance in the movies across frames and pixels - See Supplementary Materials), in order to assess whether this quantity could have confounded the estimate of RSA with the arousal and aroval model. We anticipate that the nature of the results is virtually identical with or without regressing motion energy, therefore we provide in the Results section those obtained with the simpler model.

## Results

### Behavioural data

#### Congruence between the emotion ratings across participants

Participants were instructed to rate each movie separately on the intensity of anger, fear, pain, disgust, sadness and happiness they perceived. To examine how consistently participants rated these movies, we compared each rating vector (6 values) of a given participant and movie with the mean rating vector of all other participants (i.e. in a leave-one-out fashion) for this movie (Fig. 2). The inter-subject correlation (ISC) of the ratings was high for most participants (mean = 0.85, sd = 0.23). No participants were excluded due to low ISC, despite one participant (07) having an ISC at more than 2 sd from the mean (we anticipate that the nature and significance of the RSA results did not change if this participant was excluded from further analyses).

**Fig. 2.**
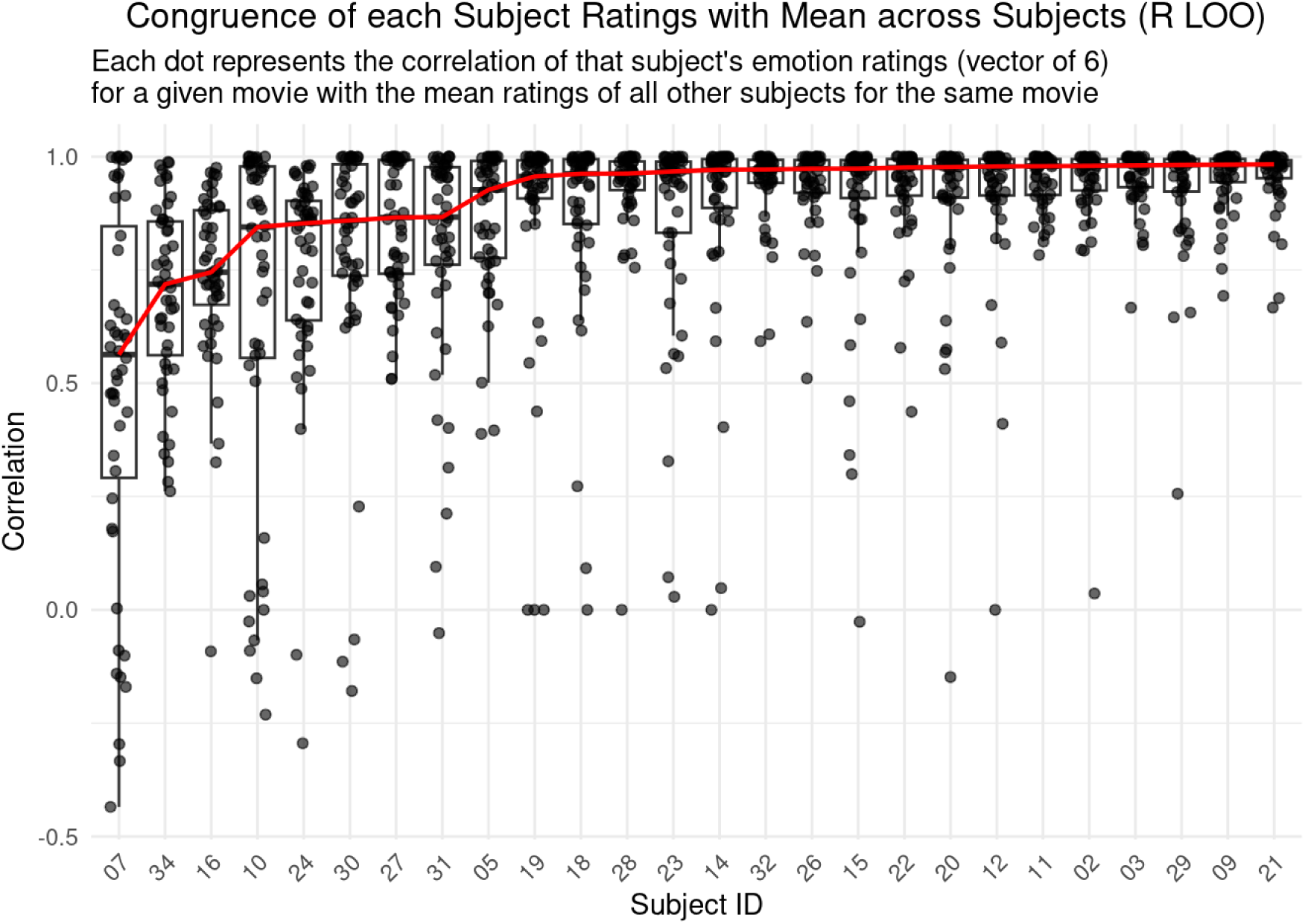
: Congruence of each subject’s ratings with the mean ratings across all other subjects. For each subject and movie, the emotion rating - a vector of 6 elements, quantifying in a scale from 0 to 10 how intensely each emotion (anger, disgust, fear, happiness, pain, sadness) was perceived in a given movie - was correlated with the mean emotion vector of all the other subjects (leave-one-out - LOO), returning 56 values (one for each movie) of congruence scores for each subject (y axis). The red line connects the median congruence across subjects. The x axis shows the different subject number arranged by increasing median congruence score across all movies for that subject.

#### Expected varibility in Arousal and Valence ratings

The initial inspection of the distribution of arousal and intensity ratings suggested a gradient of different arousal rating across emotion movies - ranging from ‘happy’ to ‘fear’, a clear separation between the valence ratings for ‘happy’ vs other movies, as well as a marked difference between movies displaying emotions at either low or high intensity (both following a similar trend of increasing arousal and valence) (Fig. 3).

**Fig 3.**
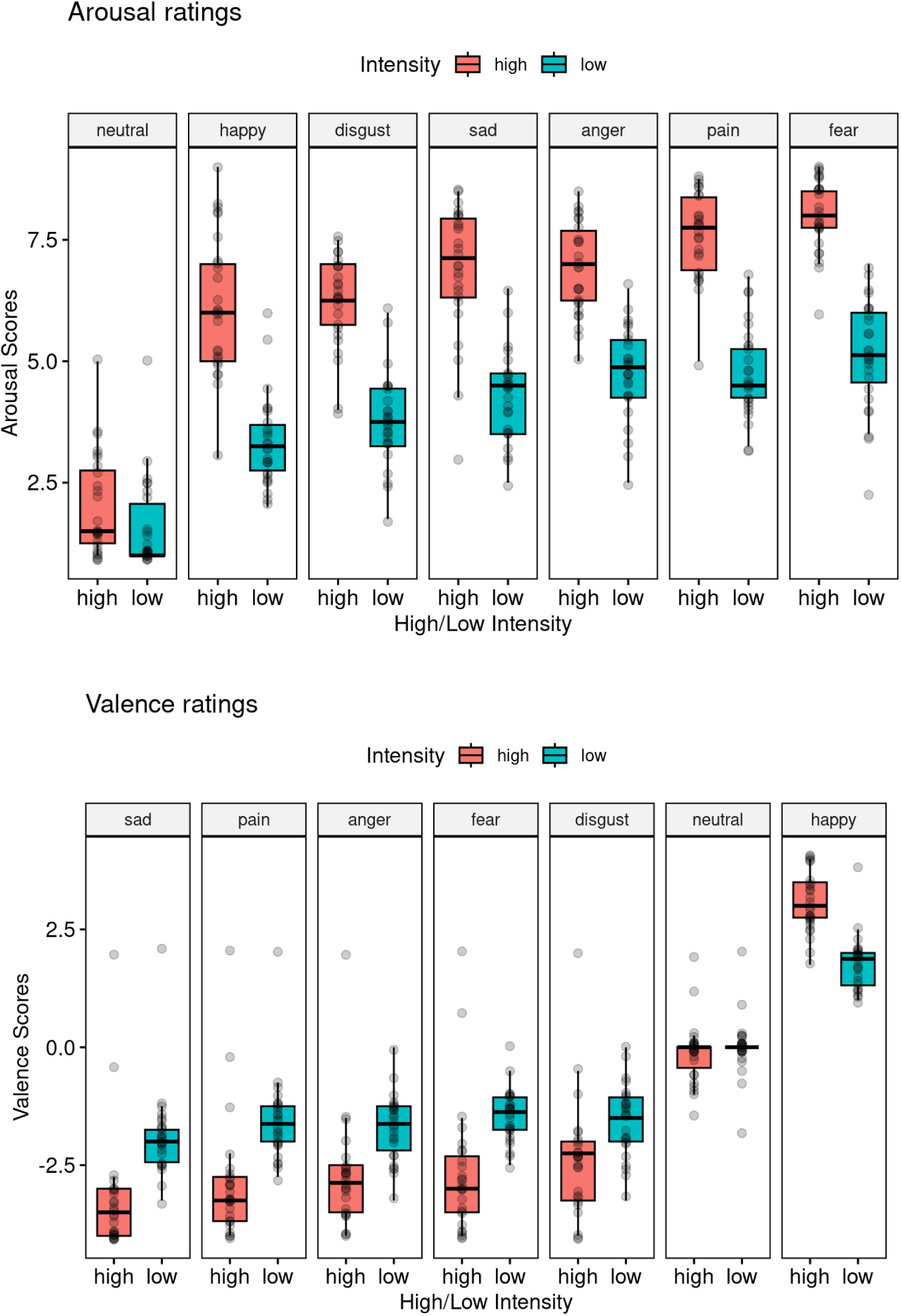
: Distribution of arousal (range 0..10) and valence (range -5..5) ratings for each movie across subjects and two levels of intensity (high and low). Each marker represents one participant’s mean rating across the 4 movies for each emotion category (or neutral) and intensity. The emotion categories have been sorted differently in the two plots to display increasing levels of perceived arousal or valence. A main significant effect of emotion as well as a significant emotion-by-intensity interaction were detected for both arousal and valence (statistics in the text). Importantly, these results also show that when the actors display a given emotion with higher intensity, this reflects on an increased perceived arousal rating, as predicted by our experimental design.

To test these observations, a repeated-measures ANOVA was conducted to examine the effects of **Emotion**, **Intensity**, and their interaction (**Emotion × Intensity**) on both Arousal and Valence ratings separately. Emotion here refers to the intended emotion displayed by the actor in the movie, not the emotional rating provided by the participants. Degrees of freedom for effects involving Emotion were corrected using the Greenhouse-Geisser method to account for violations of sphericity.

For perceived **Arousal** we detected a significant main effect of Emotion F_(3.40,84.89)_=28.56, p<.001, partial η^2^=0.533, indicating that different emotional categories had a significant effect on arousal ratings. Also the main effect of Intensity was significant, F_(1,25)_=294.84, p<.001, partial η^2^=0.922, showing that the intended intensity of the movies had a very large effect on perceived arousal. The interaction effect between Emotion and Intensity was significant, F_(4.15,103.70)_=3.40, p=.011,partial η^2^=0.120, suggesting that the effect of Emotion on arousal ratings varied depending on the level of Intensity.

Similar results were obtained for **Valence**. Also in this case both main effects of Emotion and Intensity were significant and of very large effect size: F_(2.15,53.71)_=260.85, p<.001, partial η^2^=0.913 and F_(1,25)_=41.73, p<.001, partial η^2^=0.625, respectively. The interaction effect between Emotion and Intensity was significant, F_(3.27,81.72)_=47.91, p<.001, partial η^2^=0.657.

Most of the post-hoc comparisons in either ANOVAs were also significant after Holm correction (See Supplementary Tables).

Altogether, the results of this analysis showed that movies displaying different emotions (at different intensity) resulted in different ratings along the Arousal and Valence axes, and therefore represented valid candidate stimuli to potentially elicit different brain responses along these axes.

### RSA

We tested the presence of a significant association between fmri and ratings RDMs in two large regions derived from the Yeo7 parcellation, specifically the limbic (Yeo7 ROI 5) and ventral attentional network (Yeo7 ROI 4) (Fig. 1).

Investigating the RSA in the cortical limbic regions yielded evidence of a significant association between fmri and ratings across models, as well as significant differences across models. These results were robust also for different definition of the (sub)regions of interest across atlases (Fig. 4).

**Fig. 4.**
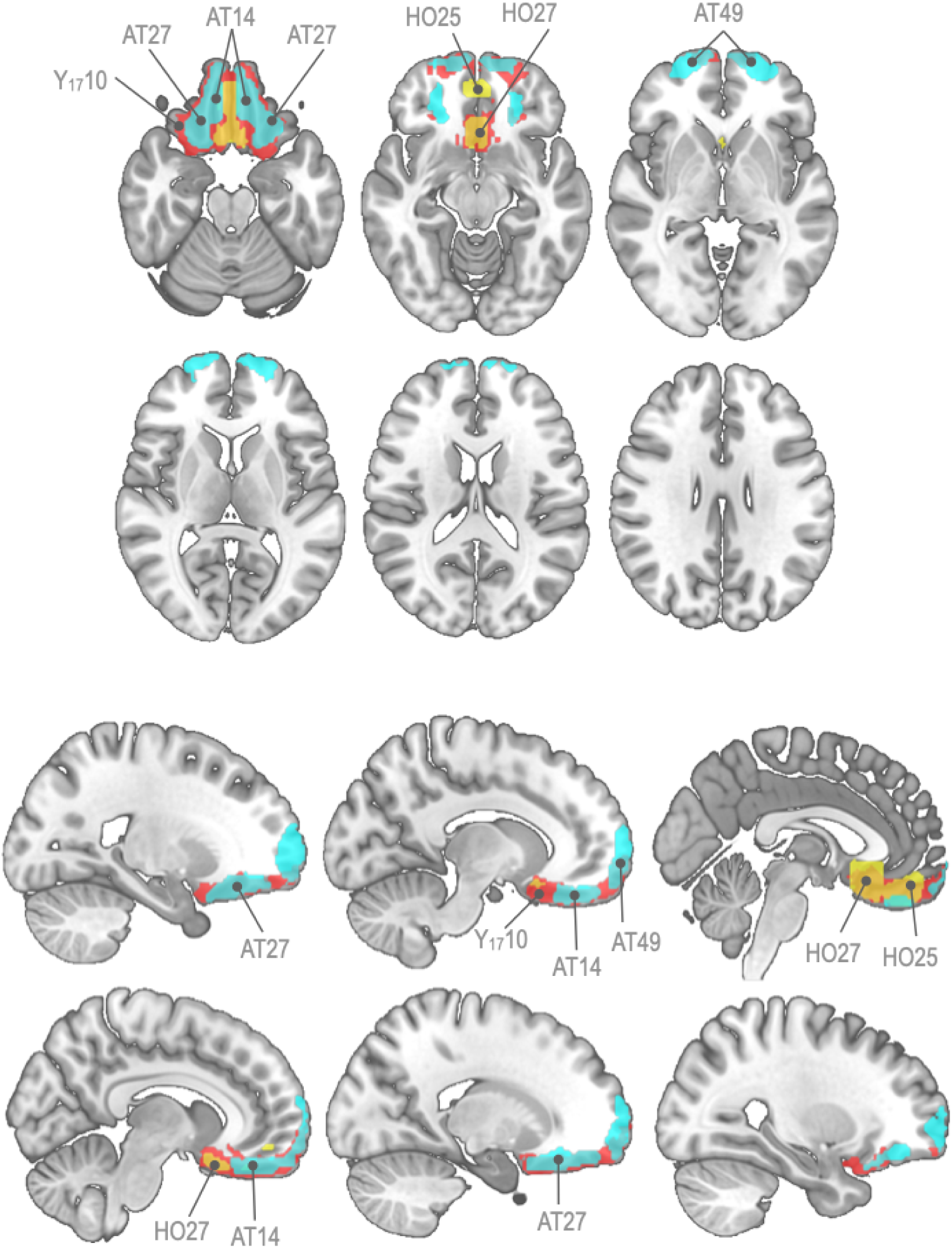
: Atlas-defined sub-regions of Yeo7 ROI #5 (Limbic) where significant differences were found in the comparison of RSA scores across models. Colors label regions in different atlases: Yeo17 subdivision (red), Harvard-Oxford atlas (HO - yellow), Anatomy Toolbox (AT - lightblue). Details about the descriptive and inferential statistics can be found in the boxplots. The p values resulting from the paired t-test comparisons within each ROI were thresholded at a significance level of p < .05 after correcting for multiple comparisons using false discovery rate. The red markers show the mean RSA value. Under each boxplot, it is shown the T value corresponding to the one-sample t-test for difference from zero and the corresponding p value (two-tailed) if smaller than p < .05. Significance markers: * = p < .05; ** = p < .01.

The initial largest region (Yeo7 ROI 5) included the lateral and medial orbitofrontal cortex (OFC), the subgenual cingulate (sgACC), the ventromedial prefrontal cortex (vmPFC), the portion of the ventral temporal lobe anterior to cerebellum and the temporal poles. This region showed highest significant RSA values for discriminating between different emotion content and to a lesser extent for rating the amount of arousal+valence in the movies. Both these models had significantly higher RSA than the arousal model. Instead the RSA scores of the latter were not significantly different from zero across participants.

This result was refined by considering separately the two subdivisions of Yeo7 ROI 5: Yeo17 ROI 10 (OFC, sgACC, vmPFC) and Yeo17 ROI 9 (temporal pole and anterior inferior temporal). The previous results for Yeo7 ROI 5 were reflected by those of Yeo17 ROI 10, while in Yeo17 ROI 9, only RSA for emotion content was significantly positive, however there was no difference between the three models.

Results obtained from the subsequent, more fine-grained atlases - the Harvard-Oxford (HO) and the Anatomy Toolbox (AT) - were consistent with the results above, in that among all HO regions sharing cortical territory with Yeo7 ROI 5, only those co-localizing with Yeo17 ROI 10 showed significant results, mirroring also in this case the results from the previous larger Yeo7/17 regions. Specifically, regions showing significant effects in the HO atlas subdivided the subgenual cortex into a caudal (HO ROI 25) and a rostral region (HO ROI 27), while the AT atlas showed significant results in four distinct regions: the medial OFC (AT ROI 14 - Fo1), the lateral OFC (AT ROI 27 - Fo3) the vmPFC (AT ROI 49 - Fp) and the Subiculum (AT ROI 82).

Across all regions, RSA scored generally higher for distinguishing different emotional content in the movies than for rating the amount of arousal + valence, while rating for arousal was always associated with RSA scores not significantly different from zero.

The other Yeo7 network we considered (Yeo7 ROI 4) encompasses the regions which have been collectively named as ‘Ventral Attentional network’ (Corbetta and Shulman 2002), which is a large-scale network encompassing regions in the posterior parietal cortex, the mid-cingulate, premotor and caudal prefrontal regions as well as the insular cortex.

When considering the whole ventral attentional network, RSA scores were significantly positive for both emotion and arousal, but with no significant difference between them. We therefore proceeded to carry out RSA scores in subregions of the Ventral Attentional network using the subregions derived from the other atlases (Harvard-Oxford and Anatomy Toolbox). The only atlas whose subregions showed an evidence of difference across tasks was the Anatomy Toolbox - which separates large insular ROIs from the Yeo and Harvard-Oxford atlas into relatively small cytoarchtectonically different subregions - and importantly only for regions in the middle and posterior insula (Fig. 5). Specifically, in the posterior ventral insula (Id5) the association between fMRI and ratings was significantly positive and different from the association with arousal+valence ratings. Instead in the adjacent Ig2 and in the middle insular Id2, arousal+valence was significantly negatively associated with fMRI parameter estimates, while there was no significant association in this region with emotion category ratings.

**Fig 5.**
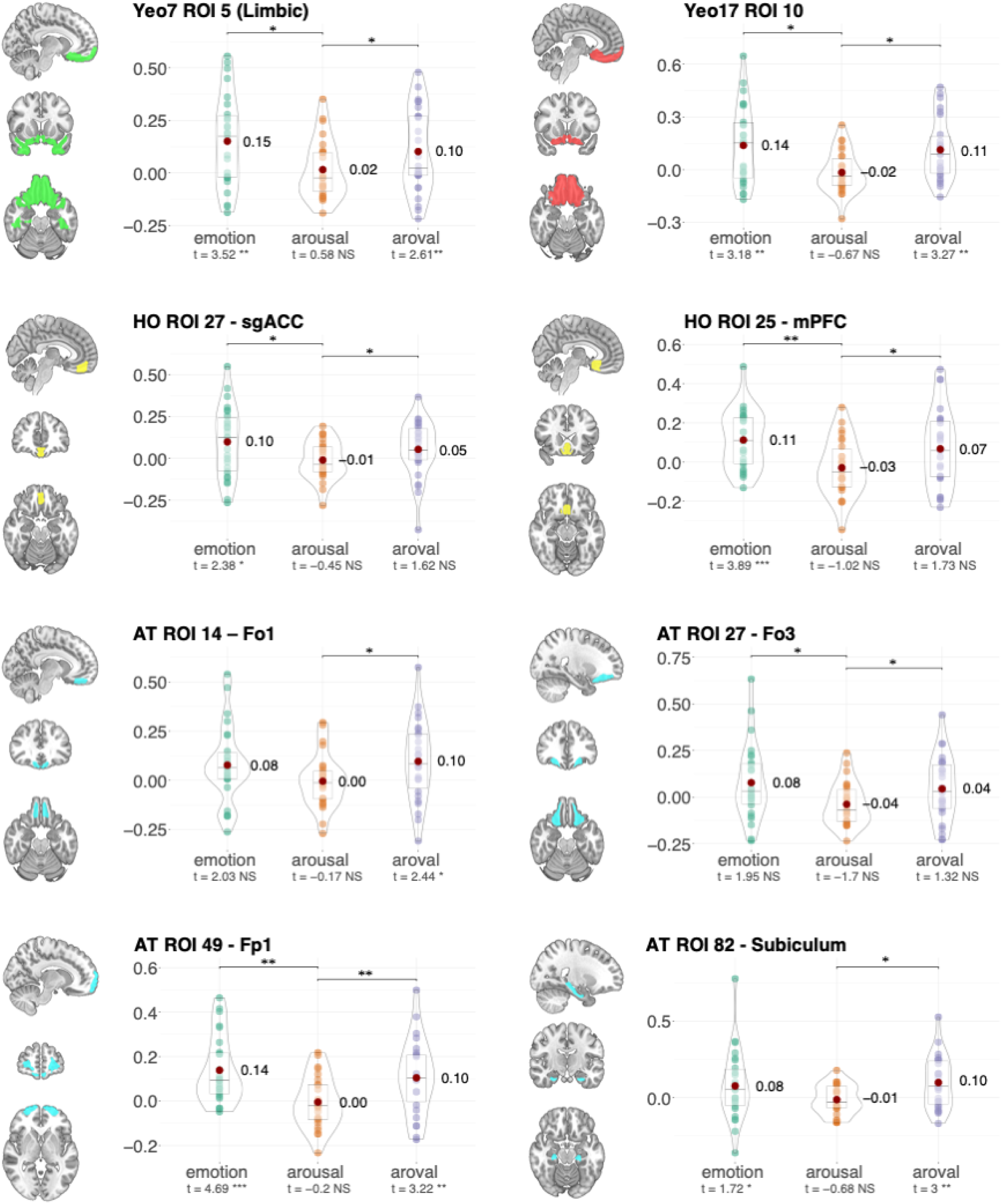
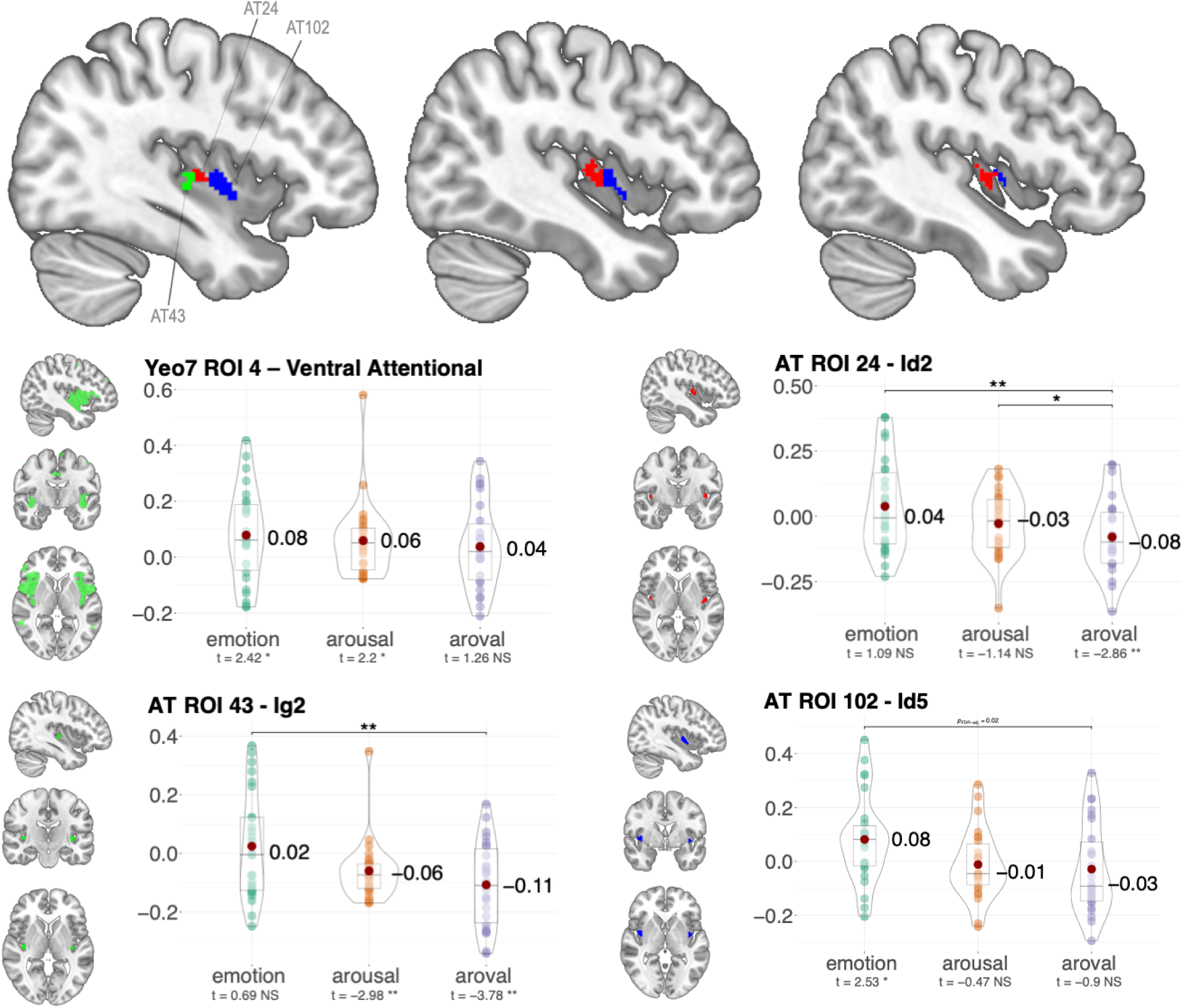
: Atlas-defined sub-regions of Yeo7 ROI #4 (Ventral Attentional) where significant differences were found in the comparison of RSA scores across models - note that no significant difference was found across models in the whole Yeo7 ROI #4 region. Colors label different ROIs within the Anatomy Toolbox (AT) atlas. No other significant differences were found considering the subregions in the Yeo17 parcellation or in the Harvard-Oxford atlas. The p values resulting from the paired t-test comparisons within each ROI were thresholded at a significance level of p < .05 after correcting for multiple comparisons using false discovery rate. The red markers show the mean RSA value. Under each boxplot, it is shown the T value corresponding to the one-sample t-test for difference from zero and the corresponding p value (two-tailed) if smaller than p < .05. Significance markers: * = p < .05; ** = p < .01.

## Discussion

After decades of research, the neural mechanisms underlying the perception of emotional facial expressions remain incompletely understood. Alongside the traditional debate between basic emotion theory - which posits discrete, evolutionarily conserved categories of emotion (Paul Ekman and Cordaro 2011; Keltner et al. 2019) - and constructivist perspectives - which emphasize the integration of core affective dimensions with contextual and cognitive factors (Barrett et al. 2019; Barrett and Wager 2006) - recent studies showed that the effect of facial emotion processing is also modulated by several contextual factors such as bodily, multi-sensory, linguistic elements and cultural factors (Saarimäki et al. 2018; Saarimäki 2021; Albohn et al. 2022; Barrett, Mesquita, and Gendron 2011; Chikazoe et al. 2014). While we acknowledge that these contextual influences play a significant role in shaping emotional perception, in the present study we specifically focused on investigating whether distinct brain regions preferentially encode categorical versus dimensional representations of emotion in response to facial expressions.

Methodologically, advances such as MultiVoxel Pattern Analysis (Haxby et al. 2001; Norman et al. 2006) and Representational Similarity Analysis (RSA; (Kriegeskorte, Mur, and Bandettini 2008)) have shifted the field from mass-univariate approaches to multivariate frameworks, enriching the characterization of the fMRI information by means of its pattern of variability across all the voxels of a given brain region. However, these approaches introduce new challenges, particularly regarding the *a priori* choice of the region of interest, which can substantially influence the results (Chang et al. 2015; Kragel and LaBar 2016) and that in turn cannot be automatically interpreted as an evidence of a multidimensional encoding (Davis et al. 2014; Hebart and Baker 2018). To address these methodological challenge, we leveraged RSA to directly compare the similarity between fMRI activity patterns elicited by emotional facial expressions and both categorical and dimensional behavioral ratings. We conducted our analyses in two different networks from the Yeo7 parcellation (Yeo et al. 2011), the first spanning the vmPFC, the OFC and the temporal poles, the other encompassing brain regions in the ventral attentional network (Corbetta and Shulman 2002; Corbetta, Patel, and Shulman 2008), anatomically centered around the insular cortex. To mitigate potential bias from initial ROIs selection, RSA was carried out for brain regions in both networks and at varying levels of anatomical granularity, thereby increasing the robustness and generalizability of our findings.

### Ventromedial prefrontal cortex

Results of the RSA analysis in the ventral prefrontal network (Yeo7 ROI 5) revealed that both the network as a whole and its constituent subregions - including vmPFC, sgACC and the frontal pole - demonstrated a significant preferential encoding of the emotion category and arousal+valence. In contrast, the was no evidence that these regions encode arousal alone. RSA scores were generally higher for the categorical model than for the arousal+valence model, however the difference did not reach significance, suggesting that these regions encode both categorical and dimensional aspects of facial emotion perception.

This pattern of findings is consistent with previous MVPA and RSA studies and meta-analyses, which showed evidence that the ventro-medial PFC is consistently included in networks that reliably encode different categories of emotion (Wager et al. 2015; Brooks et al. 2019; Roy, Shohamy, and Wager 2012; Kragel and LaBar 2016), with meta-analytic studies even suggesting a differentiation across distinct emotion categories within the subgenual cingulate (Palomero-Gallagher et al. 2015). At the same time, other studies also showed how these regions can differentiate between stimuli bearing different emotional valence (Colibazzi et al. 2010; Saarimäki et al. 2018; Chikazoe et al. 2014; Hiser and Koenigs 2018) (Yang, Tsai, and Li 2020; Sonkusare et al. 2023)(Colibazzi et al. 2010; Saarimäki et al. 2018; Chikazoe et al. 2014; Hiser and Koenigs 2018; Winker et al. 2019), with the anterior and posterior vmPFC suggesting a specialization for positive and negative emotions, respectively (Yang, Tsai, and Li 2020; Sonkusare et al. 2023).

Our results were consistent across different levels of anatomical organization, as assessed by conducting RSA both on the entire network and on its subregions, defined using resting-state connectivity (Yeo et al. 2011), anatomical landmarks (Makris et al. 2006) and average cytoarchitectonic (Eickhoff et al. 2005). While this multi-level, multi-atlas approach was designed to assess robustness across parcellations, it does not imply the existence of independent ‘sub-modules’ specialized for different encodings. Rather, the alignment of results across whole networks and subregions reinforces the view that facial emotion detection involves distributed networks, with the vmPFC as a central hub.

Beyond functional neuroimaging, this view is supported by anatomical studies in non-human primates and tract-tracing in humans, which show that the vmPFC is highly interconnected with limbic structures such as the amygdala, hippocampus, and hypothalamus, as well as with other prefrontal areas (Ongur and Price 2000; Hiser and Koenigs 2018; Haber et al. 2022; Croxson et al. 2005). These connections support the vmPFC’s role in integrating sensory, interoceptive, and reward-related information, evaluating the affective significance of stimuli, and guiding adaptive behavior. The sgACC, in particular, serves as a critical interface between the prefrontal cortex and limbic system, supporting emotion regulation and autonomic control, and is a frequent target for deep brain stimulation in mood disorders (Rudebeck et al. 2014; Johansen-Berg et al. 2008; Buchholz et al. 2024).

Clinical and lesion studies also highlight that damage to the vmPFC and associated regions often results in aberrant autonomic responses to emotionally salient stimuli, along with abnormalities in emotional introspection and regulation (Schneider and Koenigs 2017). For instance, neuroimaging studies in patients with clinical depression typically reveal hyperactivity in the subgenual cingulate and diminished connectivity between the vmPFC and the limbic system, suggesting an impaired ability to properly modulate negative emotional states. In addition, patients with lesions in the vmPFC often display maladaptive social behaviors and deficits in moral judgment, which can be attributed to disruptions in the normal encoding of emotional cues necessary for appropriate social interactions (Hiser and Koenigs 2018; Drevets, Savitz, and Trimble 2008; Wessa and Linke 2009; Tranel, Bechara, and Denburg 2002; Eslinger and Geder 2000). These clinical findings not only support the idea that this region underpins discrete categorical emotional processing but also indicate that its dysfunction can lead to an inability to modulate emotional arousal and valence appropriately.

Together with our results, these evidence from previous studies suggest that rather than a strictly dichotomous choice between categorical and dimensional aspects of the affective experience, vmPFC regions are capable of a multidimensional assessment of facial emotion perception.

### Middle and posterior insula

When RSA analysis was carried out in the Ventral Attentional Network (Yeo et al. 2011; Corbetta and Shulman 2002), we observed that at the level of the entire network, RSA scores for emotion and arousal - but not arousal+valence - were significantly different from zero, although of modest magnitude. However, when probing subregions withing this network, only cytoarchitectonically defined cortical territories in the insula (Kurth et al. 2010; Quabs et al. 2022) displayed a significant association between fMRI and ratings.

Interestingly, a distinct dissociation pattern emerged: in the posterior ventral insula (Id2, Ig2) the association with arousal+valence was significantly negative and different from the emotion category model, which failed to explain a significant amount of variance in the ratings. Conversely, region Id5 in the middle insula showed a significant positive association with emotion category ratings, but not with arousal+valence ratings. These results suggest a specialization for encoding low-level dimensional features of the facial emotion expressions in the posterior insula, while the middle insula appears to be more specialized for integrated representations of emotion categories.

This result is particularly intriguing considering the functional heterogeneity of the insula as documented by anatomical, neuroimaging and lesion studies. The posterior insula, which receives robust sensory input from the thalamus as well as parietal and occipital association cortices, is primarily involved in encoding lower-level sensory features of perceptual and interoceptive stimuli. It is sensitive to physical and homeostatic properties such as pain, temperature, and basic visceral sensations, and is often activated by the sensory intensity and unpleasantness of stimuli (Jezzini et al. 2012; Ibanez, Gleichgerrcht, and Manes 2010; Mesulam, -Marsel Mesulam, and Mufson 1985; Z. Zhang et al. 2020; Craig 2002). In contrast, more anterior sectors of the insula have been posited to serve as integrative hubs that bind these primary interoceptive signals with higher-level cognitive and contextual information, thus contributing to more abstract or integrated representations of emotion categories (Craig 2002, 2009, 2010; Uddin et al. 2017; Y. Zhang et al. 2019). For example, a recent fMRI study demonstrated that while the mid-posterior insula is active during the initial, unregulated experience of negative emotions, anterior (and to some extent middle) insula activity increases when participants engage in cognitive reappraisal, which requires abstraction and reinterpretation of emotional meaning (Z. Zhang et al. 2020). Our finding that the middle insula shows a relative preference for encoding emotion categories, when compared with the arousal+valence model, lends support to the hypothesis that this region may indeed be more sensitive to abstract, integrative features of emotion, possibly reflecting its role in the formation of coherent, categorical emotional experiences.

Clinical and lesion studies further highlight the distinct roles of insular subregions. Lesions involving the insula are known to produce disruptions in interoceptive awareness, emotional reactivity, and even in higher-order cognitive functions such as self-awareness and decision-making (Namkung, Kim, and Sawa 2017; Jones, Ward, and Critchley 2010). Notably, patients with damage to the posterior insula often exhibit deficits in processing primary somatosensory and interoceptive signals (Ibanez, Gleichgerrcht, and Manes 2010), which are crucial for evaluating basic properties of external stimuli including their arousal and valence (Berntson et al. 2011). In contrast, lesions affecting more anterior and middle portions of the insula have been associated with impairments in the subjective experience of emotion (Ibanez, Gleichgerrcht, and Manes 2010) and in the integration of complex affective information, such as trust and empathy(Belfi, Koscik, and Tranel 2015). This clinical dissociation aligns with our RSA findings, in which a higher coding of emotion categories is apparent in the middle insula (Id5), while the posterior insula (Ig2) appears more attuned to representing basic arousal and valence dimensions of emotion processing, as evidenced by its significantly negative RSA score for the arousal+valence model.

## Conclusion

Our study provides new evidence on how distinct brain networks encode the perception of emotional facial expressions, highlighting a functional diversification between the ventromedial prefrontal cortex (vmPFC) and the insula. Using Representational Similarity Analysis, we show that vmPFC regions—including the sgACC and medial OFC—encode facial emotions preferentially along categorical and valence-related dimensions, whereas the mid- to posterior insula exhibits a more nuanced profile, with posterior territories reflecting lower-level arousal/valence features and middle sectors showing sensitivity to categorical emotion content. These findings suggest that both categorical and dimensional representations coexist in the brain, but are differentially weighted across regions according to their functional roles in social and affective processing.

This distributed representational architecture helps integrating the basic emotion and constructivist perspectives on facial emotion recognition. The vmPFC appears to support higher-order evaluative processes that generate discrete, socially meaningful emotional categories, while the posterior insula encodes the bodily and sensory foundations of affective experience. The middle insula, in turn, may provide an intermediate stage of integration where interoceptive and contextual information converge into categorical representations. Taken together, our results reinforce the view that emotional empathy and recognition rely on a multilevel neural hierarchy in which categorical and dimensional representations are not mutually exclusive but dynamically interact.

From a methodological perspective, our findings underscore the value of RSA in probing representational formats beyond voxelwise activation, while also cautioning that effect sizes remain modest when stimuli differ subtly, as in facial expression paradigms. Nonetheless, the convergence of results across multiple atlases and parcellation schemes strengthens their robustness.

Clinically, our results may inform models of affective dysfunction in conditions such as depression, anxiety, and insular stroke, where disruptions in vmPFC or insular coding could compromise the balance between categorical recognition of others’ emotions and the dimensional encoding of affective states. Future work could expand this framework by examining dynamic social contexts, multimodal cues, and clinical populations, to further delineate how categorical and dimensional codes interact to support adaptive emotional understanding.

## Supplementary Materials

### fMRI parametric modulation

Before examining the effect of different emotions on the similarity of brain activity across all voxels in a brain region (RSA), we carried out a standard mass-univariate parametric modulation fMRI analysis, in which we asked how the BOLD signal elicited by observing movies of different emotions was associated with perceived emotion, valence and arousal.

To this aim, we modelled the BOLD signal at each voxel as a linear combination of the average effect of any movie plus the (de-meaned) ratings. The models for Arousal and Valence each had one predictor for rating, while the model for Emotion content had six predictors for ratings (one for each emotion).

Statistical inference was carried out by thresholding the Z images from the models at a cluster mass threshold level of Z >= 3.1 and a (corrected) cluster significance threshold of p = 0.05 (Worsley 2001).

Rating the movies on the six emotional categories or on arousal and valence content explained significant amount of variance in the BOLD activity across several regions.

Overall, Emotion and Arousal ratings significantly modulated the activity in more extensive cortical regions than Valence ratings. Brain regions whose activity was significantly modulated by any type of rating were localized in lateral extrastriate visual regions as well as in the left posterior parietal cortex (supramarginal gyrus).

Ratings for emotion and arousal showed an exclusive prominent effect in several frontal (medial and lateral) premotor regions as well as in the inferior frontal gyrus and in the adjacent right anterior dorsal insula.

Emotion ratings also significantly modulated brain activity in the mid-cingulate and in the subgenual cingulate cortex, the latter showing a significant effect in more rostral cortical territories for Arousal ratings. At the subcortical level, Emotion and Arousal ratings modulated the activity of the Putamen and of several cerebellar regions.

**Supplementary Fig. 1:**
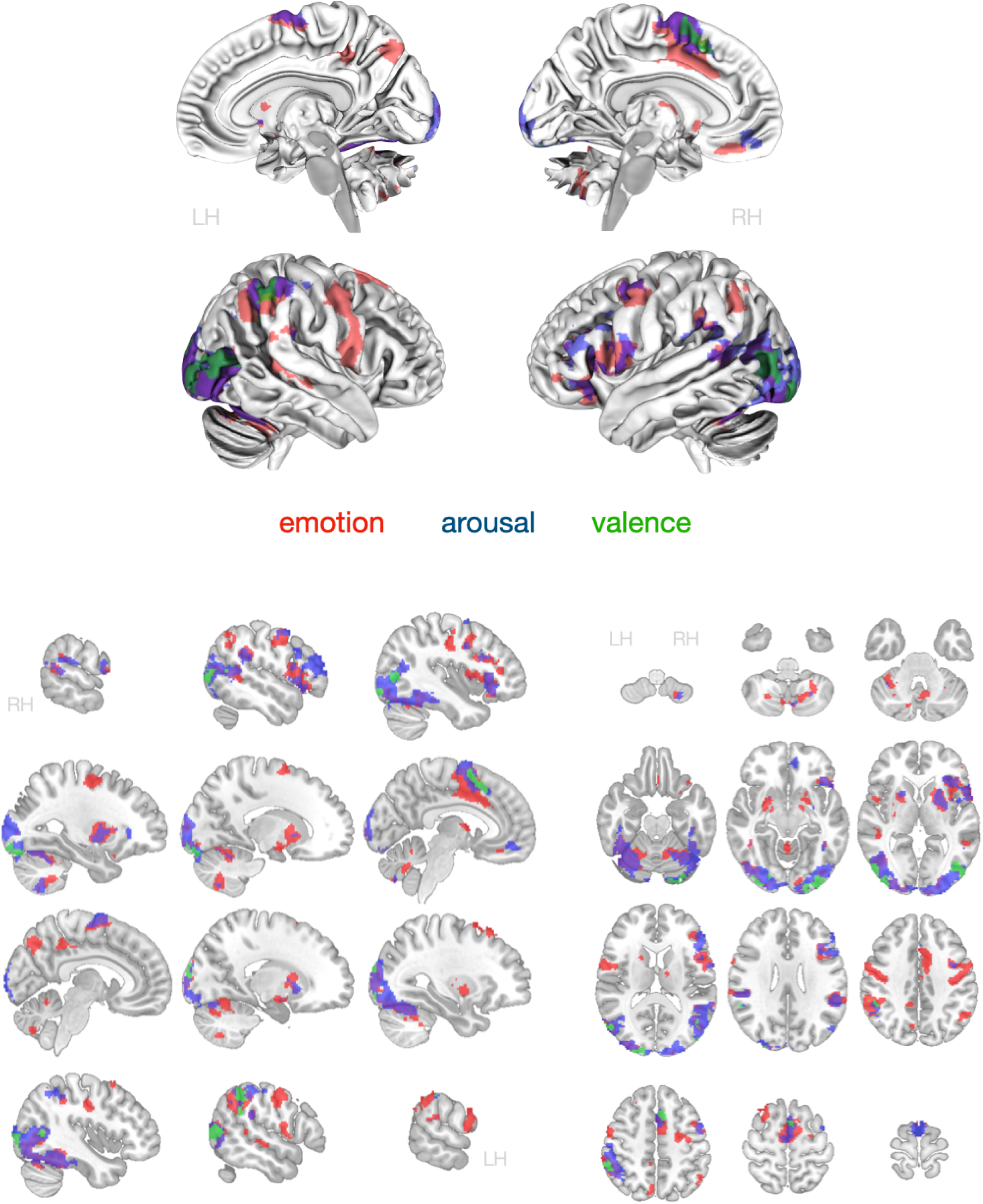
Voxelwise parametric modulation analysis, showing the effect of ratings after regressing out the main effect of movie presentation for each model: emotion, arousal, valence. Overlays represent voxels significant at a cluster mass threshold level of Z >= 3.1 and a (corrected) cluster significance threshold of p = 0.05 (Worsley 2001). Description of the significant voxels in the main text.

### ANOVA emotion-by-intensity for Arousal ratings

Supplementary tables for main and post-hoc results for the repeated-measures ANOVA examining the effects of **Emotion**, **Intensity**, and their interaction (**Emotion × Intensity**) on Arousal and Valence ratings separately.

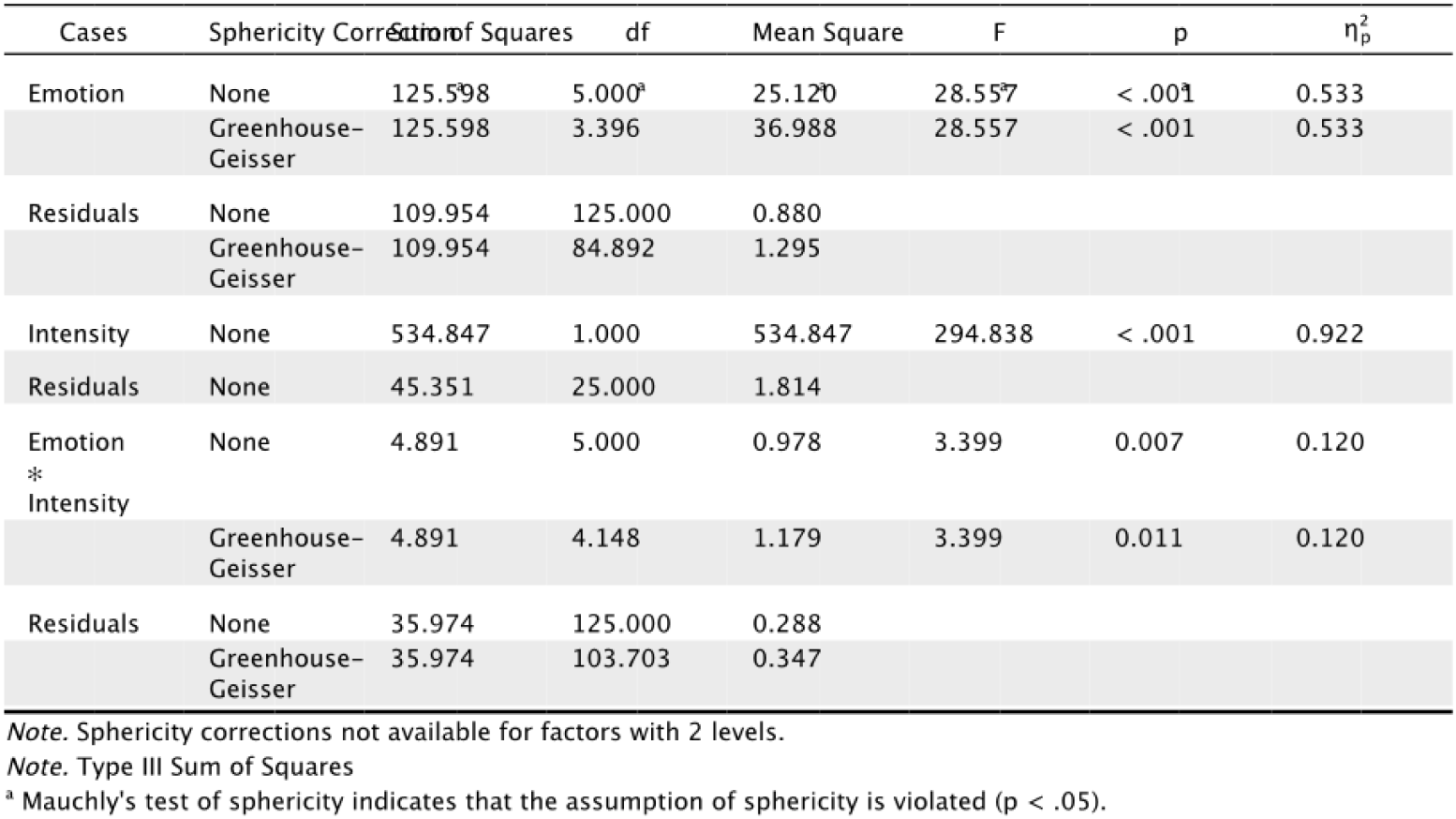

| Cases | Sphericity Correction | Sum of Squares | df | Mean Square | F | p | $\eta_p^2$ |
| --- | --- | --- | --- | --- | --- | --- | --- |
| Emotion | None | 125.598 | 5.000<sup>a</sup> | 25.120 | 28.557 | < .001 | 0.533 |
|  | Greenhouse-Geisser | 125.598 | 3.396 | 36.988 | 28.557 | < .001 | 0.533 |
| Residuals | None | 109.954 | 125.000 | 0.880 |  |  |  |
|  | Greenhouse-Geisser | 109.954 | 84.892 | 1.295 |  |  |  |
| Intensity | None | 534.847 | 1.000 | 534.847 | 294.838 | < .001 | 0.922 |
| Residuals | None | 45.351 | 25.000 | 1.814 |  |  |  |
| Emotion * Intensity | None | 4.891 | 5.000 | 0.978 | 3.399 | 0.007 | 0.120 |
|  | Greenhouse-Geisser | 4.891 | 4.148 | 1.179 | 3.399 | 0.011 | 0.120 |
| Residuals | None | 35.974 | 125.000 | 0.288 |  |  |  |
|  | Greenhouse-Geisser | 35.974 | 103.703 | 0.347 |  |  |  |

Note. Sphericity corrections not available for factors with 2 levels.
Note. Type III Sum of Squares
<sup>a</sup> Mauchly's test of sphericity indicates that the assumption of sphericity is violated ( $p < .05$ ).

**Post Hoc Tests**

*Post Hoc Comparisons-Emotion*

Post Hoc Tests Post Hoc Comparisons – Emotion
|  |  | Mean Difference | SE | t | Cohen's d | Pholm |
| --- | --- | --- | --- | --- | --- | --- |
| anger | disgust | 0.822 | 0.172 | 4.780 | 0.799 | < .001 |
|  | fear | -0.750 | 0.169 | -4.449 | -0.729 | 0.001 |
|  | happy | 1.077 | 0.198 | 5.450 | 1.047 | < .001 |
|  | pain | -0.365 | 0.175 | -2.086 | -0.355 | 0.142 |
|  | sad | 0.245 | 0.194 | 1.267 | 0.238 | 0.370 |
| disgust | fear | -1.572 | 0.159 | -9.866 | -1.528 | < .001 |
|  | happy | 0.255 | 0.187 | 1.363 | 0.248 | 0.370 |
|  | pain | -1.187 | 0.118 | -10.070 | -1.154 | < .001 |
|  | sad | -0.577 | 0.121 | -4.772 | -0.561 | < .001 |
| fear | happy | 1.827 | 0.262 | 6.975 | 1.776 | < .001 |
|  | pain | 0.385 | 0.146 | 2.630 | 0.374 | 0.058 |
|  | sad | 0.995 | 0.205 | 4.865 | 0.967 | < .001 |
| happy | pain | -1.442 | 0.202 | -7.151 | -1.402 | < .001 |
|  | sad | -0.832 | 0.225 | -3.700 | -0.808 | 0.006 |
| pain | sad | 0.611 | 0.174 | 3.502 | 0.594 | 0.009 |

Note. P-value adjusted for comparing a family of 15
Note. Results are averaged over the levels of: Intensity

*Post Hoc Comparisons-Intensity*

Post Hoc Comparisons – Intensity
|  |  | Mean Difference | SE | t | Cohen's d | Pholm |
| --- | --- | --- | --- | --- | --- | --- |
| high | low | 2.619 | 0.153 | 17.171 | 2.545 | < .001 |

Note. Results are averaged over the levels of: Emotion

### ANOVA emotion-by-intensity for Valence ratings

| Cases | Sphericity Correction | Sum of Squares | df | Mean Square | F | p | $\eta_p^2$ |
| --- | --- | --- | --- | --- | --- | --- | --- |
| Emotion | None | 938.232 | 5.000<sup>a</sup> | 187.646 | 260.854 | < .001 | 0.913 |
|  | Greenhouse-Geisser | 938.232 | 2.148 | 436.743 | 260.854 | < .001 | 0.913 |
| Residuals | None | 89.919 | 125.000 | 0.719 |  |  |  |
|  | Greenhouse-Geisser | 89.919 | 53.706 | 1.674 |  |  |  |
| Intensity | None | 43.688 | 1.000 | 43.688 | 41.732 | < .001 | 0.625 |
| Residuals | None | 26.172 | 25.000 | 1.047 |  |  |  |
| Emotion * Intensity | None | 65.578 | 5.000<sup>a</sup> | 13.116 | 47.911 | < .001 | 0.657 |
|  | Greenhouse-Geisser | 65.578 | 3.269 | 20.062 | 47.911 | < .001 | 0.657 |
| Residuals | None | 34.219 | 125.000 | 0.274 |  |  |  |
|  | Greenhouse-Geisser | 34.219 | 81.719 | 0.419 |  |  |  |

Note. Sphericity corrections not available for factors with 2 levels.
Note. Type III Sum of Squares
<sup>a</sup> Mauchly's test of sphericity indicates that the assumption of sphericity is violated ( $p < .05$ ).

*Post Hoc Comparisons - Emotion*

Post Hoc Comparisons – Emotion
|  |  | Mean Difference | SE | t | P<sub>Holm</sub> |
| --- | --- | --- | --- | --- | --- |
| anger | disgust | -0.260 | 0.096 | -2.717 | 0.071 |
|  | fear | -0.154 | 0.107 | -1.434 | 0.492 |
|  | happy | -4.639 | 0.234 | -19.797 | < .001 |
| | pain | -3.331$\times 10^{-16}$ | 0.109 | -3.050$\times 10^{-15}$ | 1.000 |
|  | sad | 0.346 | 0.118 | 2.925 | 0.051 |
| disgust | fear | 0.106 | 0.118 | 0.893 | 0.761 |
|  | happy | -4.380 | 0.227 | -19.324 | < .001 |
|  | pain | 0.260 | 0.129 | 2.010 | 0.277 |
|  | sad | 0.606 | 0.136 | 4.449 | 0.002 |
| fear | happy | -4.486 | 0.234 | -19.208 | < .001 |
|  | pain | 0.154 | 0.083 | 1.847 | 0.306 |
|  | sad | 0.500 | 0.116 | 4.317 | 0.002 |
| happy | pain | 4.639 | 0.252 | 18.442 | < .001 |
|  | sad | 4.986 | 0.257 | 19.388 | < .001 |
| pain | sad | 0.346 | 0.093 | 3.728 | 0.008 |

Note. P-value adjusted for comparing a family of 15
Note. Results are averaged over the levels of: Intensity

*Post Hoc Comparisons - Intensity*

Post Hoc Comparisons – Intensity
|  |  | Mean Difference | SE | t | P<sub>Holm</sub> |
| --- | --- | --- | --- | --- | --- |
| high | low | -0.748 | 0.116 | -6.460 | < .001 |

Note. Results are averaged over the levels of: Emotion

### TSNR

Since most of our results were located in the vmPFC and subgenual ACC, which are highly sensitive to susceptibility artifacts, in addition to the fMRI acquisition with the task, we also acquired other fMRI volumes with the same acquisition parameters and reverse phase-encoding (along the anterior-posterior axis) direction and then process them with FSL topup (Jesper L. R. Andersson, Skare, and Ashburner 2003) to correct for susceptibility-induced distortion.

We visually inspected the outcome of topup and also produce a image of the TSNR across subjects, which shows the results of the distortion correction (Suppl. Fig. 2-right).

**Supplementary Fig. 2:**
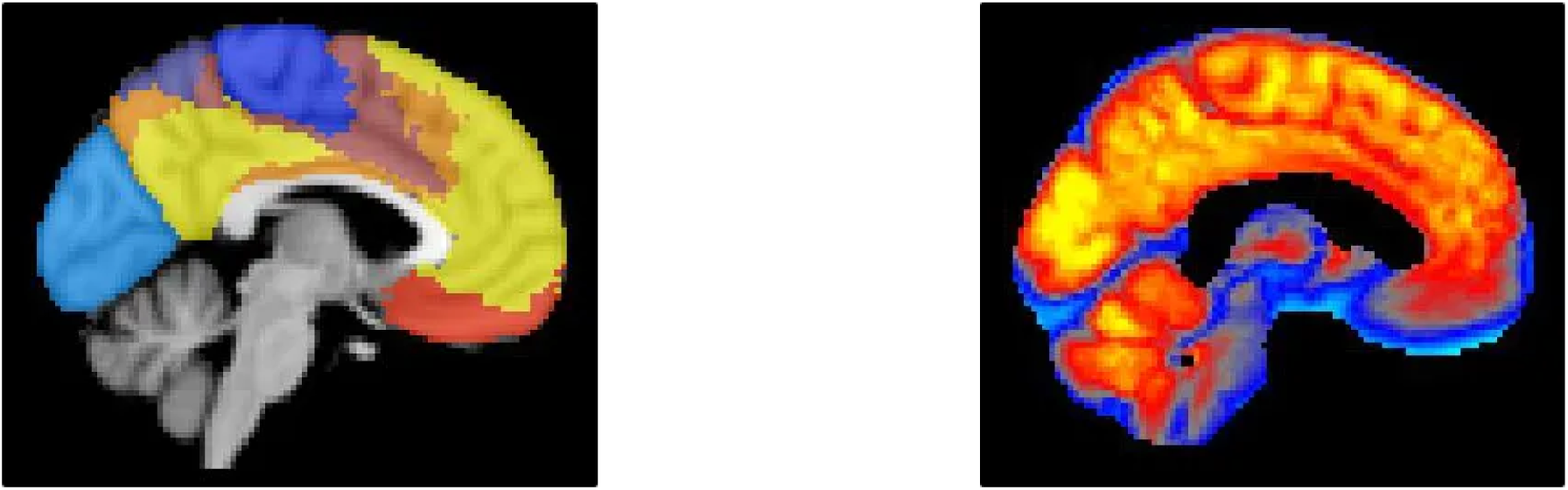
**Left:** Yeo7 regions, with ROI5 in red. **Right:** Mean TSNR across all subs in a GM prior mask. A GM prior mask was used to have an estimate of TSNR in GM, as highest value of TSNR are in WM. TSNR range: outside of the brain [blue : 5..10], sgACC [grey to red : 20..25], OCC [red to yellow: 30..35]

### Motion Energy does not affect the main results of RSA

The analysis of behavioural ratings showed that movies for different emotion categories bear different amount of perceived arousal (See Main Text “Expected variability in Arousal and Valence ratings” and relative figures). The different amount of perceived arousal, however, might not be only due to the different emotion category itself, but also to other parameters of the stimulus, such as the amount of motion in the movie.

In order to examine this possibility, we analysed differences in motion energy across all movies. Motion energy was calcuated as the sum of variability - across frames - for every pixel of the movie. Using this procedure we found indeed a significant difference in motion energy across all emotion category movies, as shown in the ANOVA table below.

**Supplementary Fig. 3:**
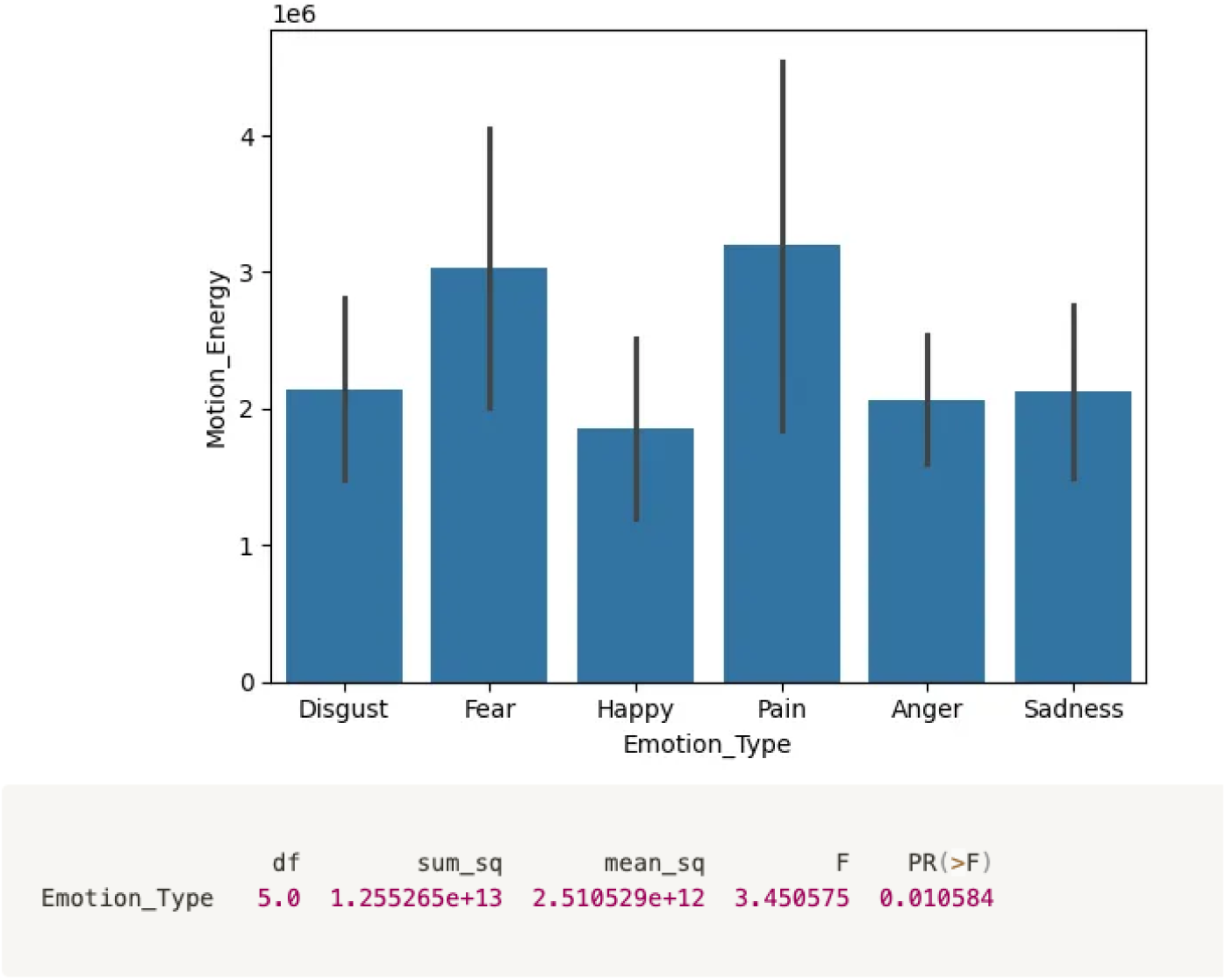
Boxplot (mean ± std) of motion energy across movies.

In order to assess whether motion energy was specifically related to the arousal ratings, we then proceeded to calculate the RDM (Representation Dissimilarity Matrix - See Main Text) for motion energy just like we did for all other behavioural ratings. Finally, we estimated the mean correlation - across participants - betwen motion energy and movie ratings (arousal, valence, aroval, emotion category) off-diagonal values in their respective RDMs (Suppl. Fig. 4). This analysis revealed that motion energy explained on average about 33% of the variance in arousal ratings RDM, and 9% of the variance in the arousal+valence (aroval) RDM. This estimate was consistent when calculating motion energy as a sum across the variances of each pixel in the video (Motion energy scalar) as well as when motion energy was calculated as a vector of values quantifying motion energy for each pixel in a downsampled version of the video (Motion energy vector).

**Supplementary Fig. 4:**
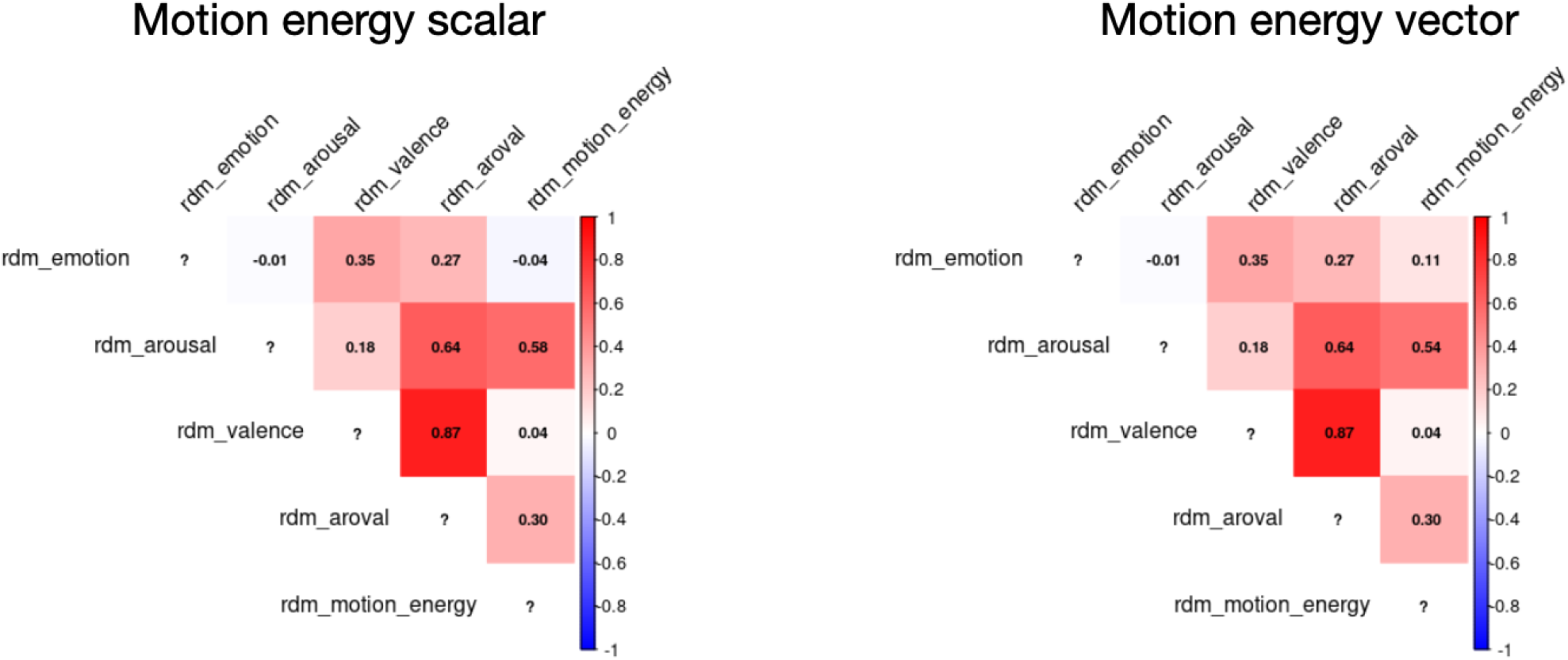
Correlation between the (off-diagonal values of the) RDM for motion energy and for ratings. The RDM for motion energy was calculated for each movie either considering the whole movie frame (left) - which returns one motion energy value for each movie - or a subsampled version of the movie (right) - where each movie returns a vector of motion energy values, one for each pixel in the subsampled movie. The results (right-most column of the matrix) were virtually identical in either versions of this calculation.

In order to estimate the effect of motion energy on our RSA analysis, we then repeated the RSA after regressing out motion energy from the ratings’ RDM. Notably, the nature of the RSA results remained virtually invariate in the two versions, with the only difference of medial subgenual cingulate and ventromedial OFC as defined by the Harvard-Oxford atlas. These result suggest that the RSA between fMRI and perceived arousal captures a different compartment of variability which is not primarily explainable in terms of the motion energy in the movies. For this reason, we still present in the Main Text the results from the RSA analysis obtained with the simpler model, that is without regressing out motion energy.

